# Recombinant inbred line panels inform the genetic architecture and interactions of adaptive traits in *Drosophila melanogaster*

**DOI:** 10.1101/2024.05.14.594228

**Authors:** Tiago da Silva Ribeiro, Matthew J. Lollar, Quentin D. Sprengelmeyer, Yuheng Huang, Derek M. Benson, Megan S. Orr, Zachary C. Johnson, Russell B. Corbett-Detig, John E. Pool

**Affiliations:** Laboratory of Genetics, University of Wisconsin-Madison, Madison, WI, 53706, USA; Department of Integrative Biology, University of Wisconsin-Madison, Madison, WI, 53706, USA; Genomics Institute, University of California, Santa Cruz, Santa Cruz, CA, 95064, USA; Department of Biomolecular Engineering, University of California, Santa Cruz, Santa Cruz, CA, 95064, USA

**Keywords:** epistasis, Quantitative Trait Loci (QTL), adaptation, quantitative genetics, *Drosophila melanogaster*

## Abstract

The distribution of allelic effects on traits, along with their gene-by-gene and gene-by-environment interactions, contributes to the phenotypes available for selection and the trajectories of adaptive variants. Nonetheless, uncertainty persists regarding the effect sizes underlying adaptations and the importance of genetic interactions. Herein, we aimed to investigate the genetic architecture and the epistatic and environmental interactions involving loci that contribute to multiple adaptive traits using two new panels of *Drosophila melanogaster* recombinant inbred lines (RILs). To better fit our data, we re-implemented functions from R/qtl (Broman *et al*. 2003) using additive genetic models. We found 14 quantitative trait loci (QTL) underlying melanism, wing size, song pattern, and ethanol resistance. By combining our mapping results with population genetic statistics, we identified potential new genes related to these traits. None of the detected QTLs showed clear evidence of epistasis, and our power analysis indicated that we should have seen at least one significant interaction if sign epistasis or strong positive epistasis played a pervasive role in trait evolution. In contrast, we did find roles for gene-by-environment interactions involving pigmentation traits. Overall, our data suggest that the genetic architecture of adaptive traits often involves alleles of detectable effect, that strong epistasis does not always play a role in adaptation, and that environmental interactions can modulate the effect size of adaptive alleles.

## Introduction

Adaptive evolution is the result of natural selection, which acts by increasing the frequency of beneficial traits in a population. A number of important uncertainties persist regarding the genetic architectures of adaptive trait changes in nature, including the number and effect sizes of contributing variants, and their frequencies before and after the action of positive selection (Pritchard *et al*. 2010, Savoleinen *et al*. 2013, Barghi *et al*. 2020). Furthermore, research on the genetic basis of adaptation has largely been centered on the individual additive effects of each gene underlying the adaptive trait, whereas the role of gene-by-gene and gene-by-environment interactions in adaptation has remained more elusive (Whitlock *et al*. 1995, Malmberg & Mauricio 2005, Martin & Lenormand 2006, Bank 2022).

Theoretical and empirical studies have yielded varied findings concerning the number and strength of loci contributing to adaptive trait change. Fisher (1930) argued that traits were governed by numerous genetic variants of very small magnitude, and that natural selection proceeded via minute incremental changes in allele frequency at these loci. Extending Fisher’s geometric model, Orr (1998) found that in a sequential model of genetic adaptation, a few larger effect sizes should tend to be followed by many smaller ones, yielding an exponential distribution of effects. However, adaptation in natural populations may sometimes depart from the assumptions of such models in multiple respects. First, the role of standing variation (as opposed to new mutations) in the adaptive process continues to be investigated (Fuhrmann *et al*. 2023, Schlötterer 2023), and these two sources of genetic variation may differ in their effect sizes, dominance, and other properties (e.g. Orr & Betancourt 2001, Ségurel & Bon 2017, Bomblies & Peichel 2022). Second, the genetic architecture of local adaptation, in which populations in a heterogeneous landscape adapt to local conditions in the presence of migration, has a stronger bias towards alleles of large effect sizes (Griswold 2006, Yeaman & Whitlock 2011, Yeaman & Otto 2011). Third, interactions among genes impacting phenotypes (*i.e.* epistasis) may alter the dynamics of adaptive variants, as further discussed below.

In regards to the role of epistasis in adaptation, a discussion has existed since the onset of the Modern Synthesis, with some arguing that it is largely irrelevant to the trajectory of Darwinian selection (Fisher 1930, Cohan *et al*. 1989, Hill *et al*. 2008, Crow 2010) and others proposing that it plays an important role (Fenster *et al*. 1997, Hansen 2013, Wright 1931). The above literature suggests that on the one hand, epistasis might not be expected to strongly affect adaptation because alleles at unlinked loci can segregate independently and selection would act on the combined effect of each allele against different genetic backgrounds; and yet, epistasis has the potential to alter the rate at which adaptive trait changes can occur by producing larger or smaller effects in some of these backgrounds, and in certain contexts, it may wield a stronger influence on the course of adaptation (Cohan *et al*. 1989, Crow 2010, Fenster *et al*. 1997, Fisher 1930, Hansen 2013, Hill *et al*. 2008, Wright 1931).

Epistasis has been shown to play a role in many evolutionary processes, including the evolution of sex and recombination, speciation, canalization, maintenance of genetic variation, and the evolution of co-adapted gene complexes (Hansen 2013). How epistatic interactions alter the response to directional selection is determined by the nature of the epistatic interaction (Hansen 2013). Some kinds of epistasis affect the magnitude of the phenotypic effects, such as positive and negative epistasis, and another kind, called sign epistasis, changes the direction of the effect. Multiple theoretical and simulation studies have investigated how each of these epistatic scenarios might influence the adaptive process (Carter *et al*. 2005; Hansen *et al*. 2006; Yukilevich *et al*. 2008; Barton 2017). For positive (synergistic) epistasis, in which the combined effect of both alleles is greater than the sum of the individual effects, additive genetic variance would increase, and the mean fitness in an adapting population is expected to change faster and reach a higher plateau than without epistasis. For negative epistasis, in which the combined effect is lower than the sum of the individual effects but still qualitatively consistent with the additive effects, additive genetic variance decreases and the mean fitness during adaptation changes more slowly and reaches a lower plateau than without epistasis. Particularly in the context of adaptation, negative epistasis is also referred to as diminishing returns epistasis. For sign epistasis, whether the allele will contribute to a larger or smaller phenotype will depend on the allele at another locus. In this case, the evolutionary dynamics can be more complex, and the fate of one allele is attached to the other, but while both variants persist, their interaction may impede adaptation. Furthermore, to the extent that epistasis masks the phenotypic influence of some variants, it may facilitate the accumulation of cryptic genetic variation for a given trait, which may subsequently be exposed to selection when elements of the genetic background (or the environment) change (Gibson & Dworkin 2004, Steiner *et al*. 2007).

Despite the pervasiveness of epistasis at the genome-wide scale (Huang *et al*. 2012), as well as between adaptive variants within proteins (Lunzer *et al*. 2010, Gong & Bloom 2014, Starr & Thorthon 2016), the prevalence of epistatic interactions among adaptive variants that contribute to the evolution of more complex adaptive traits remains incompletely understood. Studies of crop domestication have reported examples of epistasis among artificially selected alleles, including grapevine varieties (Duchêne *et al*. 2012) and many examples in maize domestication (Stitzer & Ross-Ibarra 2018). Still, compared to wild alleles, domesticated alleles were usually less sensitive to the genetic background (reviewed in Doust *et al*. 2014). Epistasis was also found among adaptive quantitative trait loci (QTLs) underlying flowering time in *Arabidopsis thaliana* (Juenger *et al*. 2002). In a recent genome-wide screen in yeast, using genes known to be involved in a high number of interactions, 24% of the adaptive variants were strain-specific and indicative of epistasis (Ang *et al*. 2023). Positive epistasis has been shown to underlie the evolution of bacteria with multiple antibiotic resistance alleles (Trindade *et al*. 2009). Examples in animals involve epistasis between alleles underlying adaptive coloration in butterflies (Papa *et al*. 2013), negative epistasis for traits underlying fitness in populations of *Drosophila melanogaster* experimentally selected on a chronic larval malnutrition regimen (Vijendravarma & Kawecki 2013), and epistasis between genes underlying coat color in oldfield mice, in which the effect of one allele is only expressed in the presence of another allele (Steiner *et al*. 2007).

The genetic architecture of adaptive traits can also be affected by gene-by-environment interactions because the effect of a given allele can change depending on its environment (Zeng *et al*. 1999, Mackay 2001). Transcriptome analysis of *D. melanogaster* reared at 20 different environments have shown that expression of ∼15% of genes changes detectably with environment (Zhou et al. 2012). In some cases, gene-by-environment interactions can be seen at the gene expression level but not the metabolic level (Reed et al. 2014), highlighting the complexity of interpreting its role in adaptation. Examples of interactions between the environment and adaptive alleles include *D. melanogaster* reproductive performance at different temperatures (Fry *et al*. 1998), drought stress adaptation in wheat (Mathews *et al*. 2008), and local adaptation reflected in biomass for switchgrass (Lowry *et al*. 2019). Similarly to epistatic interaction, gene-by-environment interaction can also uncover cryptic genetic variation (Gibson & Dworkin 2004). The chaperone protein Hsp90 offers an example of how genetic and environmental interactions can uncover cryptic variation underlying discrete and continuous traits (Rutherford & Lindquist 1998, McGuigan & Sgro 2009, Flynn *et al*. 2020).

Here, we contribute to the understanding of the genetic architecture of adaptive traits and the emerging knowledge about genetic and environmental interactions by identifying adaptive quantitative trait loci (QTLs) underlying recent local adaptation in natural populations of *D. melanogaster* and asking whether there is evidence of epistasis among them and, for a subset, whether they interact with the environment. Using two newly-described panels of recombinant inbred lines (RILs), we investigate several traits that appear to have evolved directly or indirectly under adaptive differentiation between an ancestral range population (Zambia) and a population from either France or highland Ethiopia, regions colonized by *D. melanogaster* approximately 2 kya (Sprengelmeyer *et al*. 2020).

The traits examined here (cold tolerance, ethanol resistance, pigmentation, song pattern, and wing length) show evidence of being either direct targets of local adaptation or else pleiotropic readouts of local adaptation targeting correlated traits. The pigmentation traits show strong correlations with environmental variables, especially ultraviolet radiation (Bastide *et al*. 2014), which reaches particularly high levels in the Ethiopian highlands. Larger wings may help Ethiopian flies navigate in cooler, thinner air, and wing size was found to show unusually strong quantitative trait differentiation (*Q_ST_* = 0.985; Lande 1992; Spitze 1993) between the Ethiopia and Zambia populations, compared to genome-wide genetic differentiation (Lack *et al*. 2016), indicating a role for selection in this trait’s differentiation. France and Zambia have markedly different ethanol resistance, also exceeding patterns of genome-wide genetic differentiation (Sprengelmeyer *et al*. 2021). Cold developmental survival shows a similarly consistent differentiation between France and Zambia in particular (Huang *et al*. 2021), with clear presumptive adaptive value in light of climate differences between these regions. A male song trait – the proportion of pulses classified as slow (Clemens *et al*. 2018) – was included based on strong population differentiation, with a *Q_ST_* of 0.797 exceeding the *F_ST_* of >96% of non-singleton SNPs (Lollar *et al*. 2025). Although directional selection has not been previously reported to have acted directly on song traits within *D. melanogaster*, it is also possible that this trait difference reflects selection on a pleiotropically connected trait, such as synaptic function in a novel thermal environment (Pool *et al*. 2017).

We also note that for the pigmentation, size, and ethanol resistance traits examined here, bulk mapping studies found that all of these population trait changes had very different genetic architectures from one mapping cross to the next (Bastide *et al*. 2016; Sprengelmeyer *et al*. 2021; Sprengelmeyer *et al*. 2022). Given that the differences included QTLs with effect size high enough to be detected across all the mapping crosses (Pool 2016), these results imply considerable segregating variation in genetic basis of each trait difference within one or both of the populations studied. However, it is possible (though as yet untested) that epistasis may also play a role in yielding distinct QTLs in different mapping crosses for these traits.

For each of the above traits, we first identify QTLs and candidate genes that may underlie each of these traits. Then, we quantify the strength of evidence for epistasis impacting each detected QTL, and then assess the overall signal of epistasis across traits. Lastly, we investigate the reaction norm of the QTLs underlying a subset of the traits, related to pigmentation, that were studied in two different temperature treatments. These investigations provide new insights into both the genetic basis of adaptive trait changes and the dependence of adaptive alleles on genetic interactions.

## Methods

### Mapping cross design

We report two new Recombinant Inbred Line (RIL) mapping panels of *Drosophila melanogaster*. Each RIL set was derived from a cross between two inversion-free inbred lines from distinct geographic populations, in each case pairing a strain from the southern-central African ancestral range of the species (Sprengelmeyer *et al*. 2020) with a strain from a cooler derived environment. One cross was between a Zambian inbred line (ZI215N, BioProject PRJNA861300) and a highland Ethiopian inbred line (EF43N, sequenced in Lack *et al*. 2016); the other was between a different Zambian inbred line (ZI418N, sequenced in Sprengelmeyer & Pool 2021) and a French inbred line (FR320N, sequenced in Lack *et al*. 2016). Each cross was allowed to interbreed in an intercross design for 13 and 12 non-overlapping generations, in the Ethiopian and French cross, respectively. Then, the offspring of individual females were inbred for 5 generations to create each RIL. We obtained 293 and 328 RILs for the Ethiopian and French panels, respectively (BioProject PRJNA861300). The Zambian lines were collected in Siavonga, the Ethiopian line was collected in Fiche, and the French line was collected in Lyon (Lack *et al*. 2015). The food used to rear the flies during the experiment and during the phenotyping assays was prepared in batches of 4.5 L of water, 500 mL of cornmeal, 500 mL of molasses, 200 mL of yeast, 54 g of agar, 45 mL of 10% tegosept (in 95% ethanol), and 20 mL of propionic acid.

### Phenotypes

Ethiopian flies have increased wing area, even after correcting size relative to their larger body mass (Lack *et al*. 2016). We measured the wing length of female flies that were at least 3-days old. 10 female flies per RIL were photographed using an Amscope SM-4TZZ-144A dissection microscope under CO_2_ anesthesia. The pictures were scored with ImageJ (Abramoff *et al*. 2004). Wing length was measured as the distance from the intersection of the L4 longitudinal vein and the anterior cross vein to the L3 longitudinal vein intersection with the wing margin.

Ethiopian flies are darker than Zambian flies (Bastide *et al*. 2014). We photographed and measured two to five females per RIL using a dissecting scope. Resolution (3584 x 2748 pixels), gain (3.0), exposure time (99.84 ms), magnification (20x), and illumination level were kept constant using an Amscope adaptor for LED lamp at maximum lighting. White balance was set at the same resolution using ColorChecker white balance. We measured three common traits for pigmentation: the pigmentation of the mesopleuron in greyscale proportion ranging from 0 for white to 1 for black (herein, mesopleuron, a thorax trait), the pigmentation of the background of the fourth abdominal segment in the same greyscale proportion (herein, A4 Background), and the proportion of the fourth abdominal segment that was covered by the black stripe (herein, stripe ratio). Relatively low correlations (computed via regression) among these three traits were observed among independent isofemale strains within the Ethiopia and Zambia populations (*r* < 0.35), except for a higher correlation between mesopleural and A4 Background for Ethiopia specifically (*r* = 0.646; Bastide *et al*. 2014).

Pigmentation is also plastically influenced by temperature, and the Ethiopian population is located in a colder region, on a high plateau 3 km above sea level. Therefore, we measured the three pigmentation traits in flies raised at two temperatures (25 °C and 15 °C), to uncover potential cryptic variation that could be present in the warmer Zambian (midpoint temperature: 25 °C) but is only expressed in the colder Ethiopian (midpoint temperature: 11 °C) population. We also report plasticity for these three pigmentation traits, as the difference in the trait measured at 25 °C and 15 °C.

French flies have higher resistance to ethanol (Cohan & Graf 1985, Sprengelmeyer & Pool 2021) and cold (Pool *et al*. 2017, Huang *et al*. 2021) than Zambian flies. Ethanol resistance was measured as the average time 300 female flies, 3- to 5-day old, from each RIL needed to become immobile when exposed to 18% ethanol. The flies were placed in 50 ml falcon tubes with ethanol-saturated tissue placed on the bottom, and the number of mobile flies was scored every 15 minutes for six hours. Cold tolerance was measured as egg to pupa viability when raised at 15 °C. Given that very few dead pupa were observed across RILs, this measure was taken as a reasonable time-saving proxy for the egg to adult viability difference previously demonstrated between the parental strains (Huang *et al*. 2021). To obtain this metric, six females and six males (3∼5-day old) were collected from each RIL and allowed to lay eggs for 15 hours. The egg to pupa viability was estimated as the number of emerged adults plus left pupa in a vial at Day 34∼35 divided by the number of eggs present in the vial after the parents were removed (Day 1).

The song pattern trait was the ratio of slow pulses over all pulses (slow and fast) in the song made by males during courtship. Although not expected to be under local environmental selection *a priori*, male courtship is often under strong sexual selection. In addition, a large number of nervous system genes appear to have evolved between warm- and cold-adapted *D. melanogaster* populations (Pool *et al*. 2017), representing one potential source of pleiotropic effects on song traits. We found that Zambian male flies display a greater fraction of slow to fast pulses than French flies (Lollar *et al*. 2025), a novel pulse mode classification recently discovered within the *D. melanogaster* species complex (Clemens *et al*. 2018). The song data was recorded in the Stern Lab at Janelia Farms Research Campus, using materials and methods described by Arthur and colleagues (Arthur *et al*. 2013). Song was annotated without human intervention with SongExplorer (Arthur *et al*. 2021). Annotations were classified with locally modified versions of BatchSongAnalysis (https://github.com/dstern/BatchSongAnalysis), using previously trained *D. melanogaster* models provided by Ben Arthur (Lollar *et al*. 2025).

We expected that there might be some level of correlation among the traits of any given cross purely due to either linkage among causative variants or else pleiotropy. We calculated the pairwise correlation between all the traits measured for each cross to investigate the degree to which they are correlated.

### DNA Extraction and Sequencing

For each RIL, we extracted DNA from five to ten female flies from each line. The flies were crushed and homogenized using pipette tips in strip tubes, in a solution of 100 μl of lysis buffer (50 mM Tris-HCl with pH = 8, 100 mM EDTA, 100 mM NaCl, and 0.5% SDS) and 2 μl of proteinase K (IBI Scientific). The samples were incubated to inactivate the proteinase K (55 °C for 5 minutes, 95 °C for 10 minutes, and 25 °C hold). The supernatant of the fly lysate was transferred to new tubes, and we added SPRI beads (Sera-mag SpeedBeads SPRI beads, [GE Healthcare 24152105050250]) in a 1.8 to 1 proportion of beads to lysate. The beads were used in a solution of PEG 8000 (9 g), Tween 20 (27.5 μl), 5M NaCl, and 3 mL TE Buffer (composed of 100 μl of 1 mM EDTA, 500 μL of 10 mM Tris HCl). 1 mL of the beads were added to the buffer mix and filled with water until the final volume was mL. The tubes were incubated for at least three minutes at room temperature for DNA to bind to the beads and then placed on a magnetic plate for five minutes to remove the beads from the solution. The supernatant was discarded, and the beads were washed with two rounds of 80% ethanol (90 μL per well). DNA was eluted beads with 30 μL of H_2_O and then used the magnetic plate to transfer only the resuspended DNA to new tubes.

Genomic libraries were prepared following Adams *et al*. (2020), with size selection, cutoff at 300 bp, and cleanup performed using the Zymo Select-a-Size DNA clean & Concentrator [catalog No. D4080]. Libraries were sequenced at the UW-Madison Biotechnology Center on the Illumina NovaSeq 6000 platform, with 150 bp paired-end reads.

### DNA Alignment and Ancestry Calling

The reads were mapped to the *D. melanogaster* (v5.57) reference genome using BWA-MEM, version 0.7.17 (Li 2013). We used Ancestry HMM (Corbett-Detig & Nielsen 2017) to estimate ancestry along the genome of each RIL, allowing for diploid ancestry calls (*i.e.* homozygous for either parent, or heterozygous). Ancestries were summarized in non-overlapping windows defined to contain 1,000 SNPs in the Zambian population (19 kb average size), using the majority ancestry call in the case of an ancestry breakpoint within the window. Window size was chosen to ensure ample variation for ancestry calling within each window, and to moderate the number of QTL and epistasis tests performed in order to ensure computational tractability, while still allowing a genomic resolution much narrower than the expected megabase scale ancestry tracts. When two adjacent windows had the same ancestry genotype across all RILs, the two windows were merged. We used a recombination map based on Comeron *et al*. (2012) but modeled only one generation of recombination when using Ancestry HMM, in order to focus on conservatively-defined ancestry switches (pipeline available on https://github.com/ribeirots/RILepi).

### QTL Mapping

To test whether adaptive loci showed evidence of epistatic interactions we first used QTL mapping to identify the adaptive loci and then performed epistasis tests between the focal loci and all other loci in the genome. Each locus is a genomic window of ∼19kb, defined and genotyped as described above.

To identify loci underlying trait changes, we employed a similar approach to the R/qtl (Broman *et al*. 2003) *scanone()* function, but with an additive framework, modifying the genotypes to be coded as numeric variables instead of categorical (https://github.com/ribeirots/RILepi), using Python3 (Van Rossum & Drake 2009). The additive modification was preferred in light of the small counts of specific genotypes due to the low heterozygosity and the variable genomic ancestry skew present in our RIL panels. We fit a model of the phenotype given the ancestry genotype of each window (Trait ∼ Marker). In our model, we coded the genotypes at Q as numeric values: 0, 1, and 2 for the homozygous Zambian ancestry, heterozygous ancestry, and Ethiopian or French homozygous ancestry, respectively.

The initial QTL mapping is based on the comparison of a model with a single QTL *versus* a model without any QTL, performed repeatedly one locus at a time. For each locus, we obtained a logarithm of the odds (LOD) score as the ratio between the log-likelihoods of the model with one QTL over the model without a QTL. Because both RIL sets had average ancestral frequencies skewed against the Zambian ancestry (68% Ethiopia ancestry and 75% France ancestry), we excluded the windows with an ancestry bias of 90% or greater for either ancestry to avoid the effects of extremely uneven sample sizes on the model. The genome-wide significance of a QTL was tested using 10,000 permutations, in which trait values were randomly shuffled. From each permutation, we saved the highest LOD score obtained for any window (excluding windows with ancestry skew of 90% or greater) and empirical LOD scores from each window were compared to the null, permutation LOD score distribution. We considered the windows whose LOD score’s p-value was lower than 0.10 as putative QTLs (*i.e.* a less than 10% chance of having a QTL signal of that strength anywhere in the genome). A p-value threshold of 0.10 was chosen aiming to include a sufficient number of QTLs to be tested for epistasis. We emphasize that these QTL p-values are trait-specific, and do not attempt to quantify the likelihood of any QTL peak for any analyzed trait having a given LOD score.

Each putative QTL surrounded by windows with a lower LOD score was considered a putative QTL peak. If two putative QTL peaks were not separated by a window with a LOD score lower than the minor peak’s LOD score minus 1.5, the minor peak was removed. Finally, the remaining putative QTL peaks were filtered to remove any peak within 10 cM of a QTL peak with a higher LOD score, in order to increase the independence of QTLs used in epistasis testing. The confidence interval for these QTLs was defined as all contiguous windows with a LOD higher than the QTL’s peak LOD score minus 1.5. Heritability-based QTL effect (*h^2^*) was calculated as described in R/qtl for an intercrossing mapping design (Broman *et al*. 2003).

The power of the QTL mapping analysis was calculated based on simulated data for each RIL panel. We simulated a normally distributed phenotype (mean = 0, standard deviation = 0.75) with additive effect of the derived allele varying from 0.05 to 0.5 in 0.05 increments and without dominance. We randomly sampled genotype distributions for each simulation from the empirical RIL panels, to a total of 10,000 simulations for each additive effect and RIL panel. We calculated the LOD score and heritability-based QTL effect (h^2^) for each simulation as described above, and calculate its p-value based on the permutations of the empirical data for each RIL panel. QTL mapping power was determined for each additive effect and RIL panel as the proportion of simulations with p-value equal to or lower than 0.1–the same threshold used in our analysis.

### Candidate Genes and Population Genetics

To identify genes potentially underlying the adaptive traits, we scanned our QTL regions for signatures of local adaptation based on population genetics. We sequenced genomes from inbred lines from each of the studied populations to calculate a haplotype-based statistics: χ_MD_ (Lange & Pool 2016) and two *F_ST_* statistics (using Reynolds *et al*. 1983), *F_ST_FullWin_* (window-wide *F_ST_* calculated using all the SNPs in the window) and *F_ST_MaxSNP_* (the highest *F_ST_* value from a single SNP within the window), that have power to distinct power to detect distinct kinds of selective events (da Silva Ribeiro *et al*. 2022). We defined as outlier windows those that fell within the top 1% of windows on the same chromosome arm for any of the above three statistics. We focused on regions with recombination rates generally above 0.5 cM/Mb (Comeron *et al*. 2012) due to more localized signatures of selection: 2.3–21.4 Mb of the chromosome X, 0.5–17.5 Mb of arm 2L, 5.2–20.8 Mb of arm 2R, 0.6–17.7 Mb of arm 3L, and 6.9–26.6 Mb of arm 3R. We selected as candidates all genes that overlapped with the outlier windows as well as the first gene up- and down-stream of the outlier window, to account for instances in which the target of selection is in a regulatory region outside the gene region. We also identified other genes within these QTLs with a known or potential association with the phenotype based on FlyBase (release FB2023_05) annotation.

### Epistasis Testing

Epistasis tests were performed between each focal QTL and all of the remaining genomic windows outside its confidence interval. The epistasis test for each window pair consisted of an interaction LOD score obtained as the ratio between the log-likelihoods of the model with two QTLs and their interaction (Trait ∼ Marker1 + Marker2 + Marker1:Marker2) *versus* a model including both QTLs but no interaction (Trait ∼ Marker1 + Marker2). The interaction LOD score for each pair was compared against permutations. Permutations were specific for each QTL analyzed. We shuffled the phenotype, fixed the focal QTL (Marker1), and calculated the interaction LOD score of Marker1 against all other windows in the genome. The highest genome-wide interaction LOD score from each permutation was kept and the empirical interaction LOD scores were compared against this null, permuted interaction LOD distribution.

### Epistasis Meta-analysis

We combined the results from all the epistasis tests across mapping crosses and phenotypes to test the hypothesis of whether adaptive QTLs have epistatic interactions. We collected the set of epistasis p-values from each QTL and its most likely interactor, quantifying the probability of obtaining any interaction term between that QTL and any partner locus in the genome with an interaction LOD score as high as the one observed. We used Fisher’s combined probability test to ask if the p-values from each QTL and their most likely interacting locus reject the null hypothesis that there is no epistasis. The p-values should follow a uniform distribution if the null hypothesis is true. Therefore, we performed an additional step to investigate how many of the lowest p-values would need to be removed from the data set to obtain a median value near 0.50, in order to ask whether our cumulative data set showed any hints of epistasis across all analyzed QTLs.

### Epistasis Power Analysis

We calculated the power of our approach and our data sets to detect an interaction by simulating phenotypes with different interaction strengths. We chose one empirical QTL with intermediate sample size and estimated the additive QTL effect among the detected QTL to serve as the basis for the simulations. The QTL chosen was detected for the abdominal background pigmentation at 25 °C, it had an estimated effect size equal to 0.035, mean trait value equal to 0.3485, and a standard deviation equal to 0.0697. The observed genotype distribution of the QTL peak window was used for the simulations. Our null simulations generated phenotypes based on a normal distribution with the observed trait mean and standard deviation, and the simulated values were modified based on the trait effect and individual genotypes (1x effect for heterozygous and 2x effect for non-Zambian homozygous). We then calculated the interaction LOD score of the focal peak window against all other genome windows outside the original QTL’s confidence interval and kept the highest genome-wide interaction LOD score obtained. We repeated this process 10,000 times to obtain a null (no interaction) distribution of interaction LOD scores.

To simulate epistasis, we used the genotype distribution of the window most likely to be interacting with our focal QTL: the window with the highest empirical interaction LOD score. We modified the empirical effect of the focal QTL on the simulated phenotype based on the genotype of the interacting genomic window. If the genotype of the interacting window was Zambia homozygous, the original effect was not modified, if it was non-Zambia homozygous, the original effect was multiplied by the interacting factor *I*, and if it was heterozygous, the original effect was intermediate (*i.e.* multiplied by (1+*I*)/2). The values of *I* we simulated varied from −2 to 4, depending on the nature of the epistatic interaction being simulated. We simulated positive epistasis effects (original QTL effect increased by the presence of the epistatic allele) with *I* ranging from 1.125 to 4. For diminishing returns negative epistasis (original QTL effect decreased by the presence of the epistatic allele), *I* ranged from 0.889 (1/1.125) to 0.5. We simulated masking epistasis (the original QTL effect nullified in the presence of the epistatic allele) with *I* = 0. Lastly, for sign epistasis (the original effect changes in sign, *e.g.* instead of the non-Zambia QTL allele producing larger trait it produces smaller traits), we examined effects from - 0.5 to −2.

Each interaction simulation was compared against the null distribution and a p-value was calculated as the proportion of null simulations with an interaction LOD score higher than the simulated interacting LOD score. Power was calculated for each interaction effect *I* as the proportion of simulations that had p-value lower than 0.05.

We also calculated the power of our meta analysis based on 14 QTLs’ interaction p-values. For each interaction effect, we sampled 14 interaction LOD score p-values and calculated (1) how often the fisher combined p-value was as extreme or greater than the one we observed and (2) how often at least one p-value was lower than the lowest p-value we obtained for our empirical data.

### Reaction Norms

To investigate genotype-by-environment interactions for pigmentation traits, we combined phenotypic data from RILs raised at the two temperature treatments (15 °C and 25 °C) with the genotype of each non-overlapping QTL. To test whether the genotype-by-environment interaction was significant, we used a linear model with the phenotype as the response variable and the genotype, environment, and genotype-by-environment interaction as the explanatory variables.

## Results

### Genotype and phenotype data

Whole genome sequences for the RILs had mean depth of coverage of 3.3X (S.D. 1.96 among RIL mean depths, Table S1) and 7.58X (S.D. 9.09, Table S2) per site for EF and FR RILs, respectively, which should be more than sufficient to call population ancestry in large chromosomal blocks (Corbett-Detig & Nielsen 2017). Ancestries for both RIL sets were skewed toward lower Zambian ancestry, with averages of 67.84% Ethiopian ancestry and 77.88% French ancestry (full ancestry genotypes in Tables S3, S4). After five generations of inbreeding, a low level of parental strain heterozygosity was present in the final panel: 8.71% and 5.3% on the Ethiopian and French panels, respectively (at the level of genomic windows). The sample size varied for each trait based on the number of RILs successfully phenotyped (Table 1, S1, S2).

**Table 1.** Summary of mapped traits. Listing the mapping cross used for each trait (EF = Ethiopia, FR = France), the number of phenotyped lines for each trait, the number of QTLs, and the number of QTLs with epistatic interaction for each phenotypic trait.

| RIL panel | Trait | Sample size | # QTLs |
| --- | --- | --- | --- |
| EF | A4 Background 25 °C | 278 | 1 |
| EF | Stripe Ratio 25 °C | 278 | 1 |
| EF | Mesopleuron 25 °C | 278 | 1 |
| EF | A4 Background 15 °C | 190 | 0 |
| EF | Stripe Ratio 15 °C | 190 | 1 |
| EF | Mesopleuron 15 °C | 190 | 1 |
| EF | A4 Background (Plast.) | 178 | 0 |
| EF | Stripe Ratio (Plast.) | 178 | 0 |
| EF | Mesopleuron (Plast.) | 178 | 0 |
| EF | Wing Length | 268 | 5 |
| FR | Cold Tolerance | 304 | 0 |
| FR | Ethanol Resistance | 277 | 3 |
| FR | Song Pattern | 294 | 1 |

The distribution of the phenotypes were unimodal for all traits with varying degrees of variance and low to moderate skew (Figures S1, S2). Correlations among the analyzed traits were examined for each RIL panel, which may indicate pleiotropy or else linkage of causative alleles. Among pigmentation traits, the thoracic mesopleuron trait was strongly correlated with abdominal background: r = 0.687 at 15 °C, r = 0.800 at 25 °C, r = 0.782 for the plasticity of these traits (Table S5). Stripe width showed moderate, significant correlations with both of those traits at 25 °C (*r* = 0.460, and *r* = 0.253), but weaker correlations at 15 °C (*r* = 0.169, and *r* = - 0.085). Abdominal background showed a non-significant correlation between the two temperatures, while the other two traits showed moderate, significant correlations between temperatures. In light of the correlations observed among many pigmentation traits, overlapping QTLs for pigmentation traits were not treated as independent QTLs, and only the QTL with the highest LOD score was used in downstream analyses. No pigmentation trait was significantly correlated with wing length (Table S5), the only other trait scored among the Ethiopian RIL panel. Among traits scored from the French RIL panel, the song pattern was weakly correlated with ethanol resistance, r^2^ = 0.13 (Table S6).

### Single locus QTL mapping

We identified potential QTLs for eight out of the thirteen investigated traits (Figure 1; Table 1). We list the genome-wide distribution of LOD scores and p-values for each trait in the Table S7. We also present the phenotypic distributions observed for each genotype at the QTL peak window (Figures S3, S4) including the pulse song usage trait that is discussed in more detail by Lollar *et al*. (2025). The traits with no detected QTLs were A4 Background at 25 °C, all the three pigmentation plasticity traits, and cold tolerance (genome-wide LOD scores shown in Figure S5). The number of QTLs per trait ranged up to five, for wing length, with other traits yielding one or three QTLs (Figure 1, Table 1). Twelve of the fourteen QTLs were significant at p < 0.05, while two marginally significant QTLs - one for A4 Background at 25 °C (p = 0.0591) and one QTL for ethanol resistance (p = 0.0693) – were also considered in downstream analyses (Table 2). The effect of each QTL, based on estimated *h^2^*, varied from 0.033 to 0.285 and we had high power (>0.8) to detect QTLs of effect size 0.1 (Figure S6).

**Figure 1.**
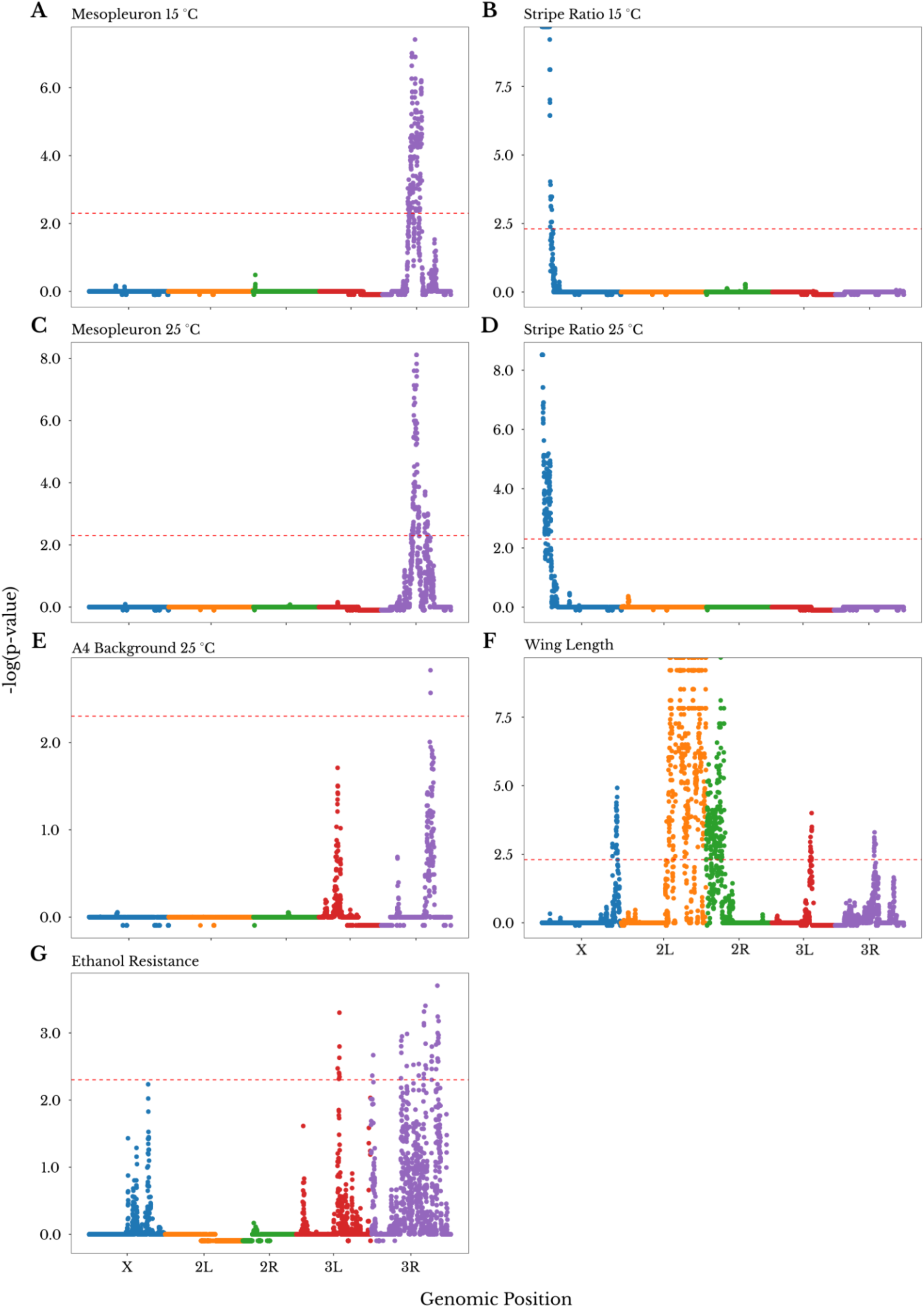
The number of QTLs for each trait ranged from 0 to 5. In this figure, only the traits with at least one identified QTL are shown (phenotype distributions for each genotype at each QTL peak window are shown in Figures S3, S4); the traits without significant QTLs (Table 1) can be seen in Figure S5. Each panel shows the −log of the (trait-specific) p-value for the LOD score of the genomic windows. A4 Background (Figure 1E) is shown here with no QTL above the threshold, but one marginally significant QTL (p-value = 0.053, Table 2). Windows filtered out for ancestry skew are given a value of −0.25. The red dashed line represents the 0.05 p-value cutoff based on 10,000 permutations. The color of the dots represents the chromosome arm of each genomic window. Note that for wing length, a single QTL spans a broad low recombination centromeric region between 2L and 2R.

**Table 2.**
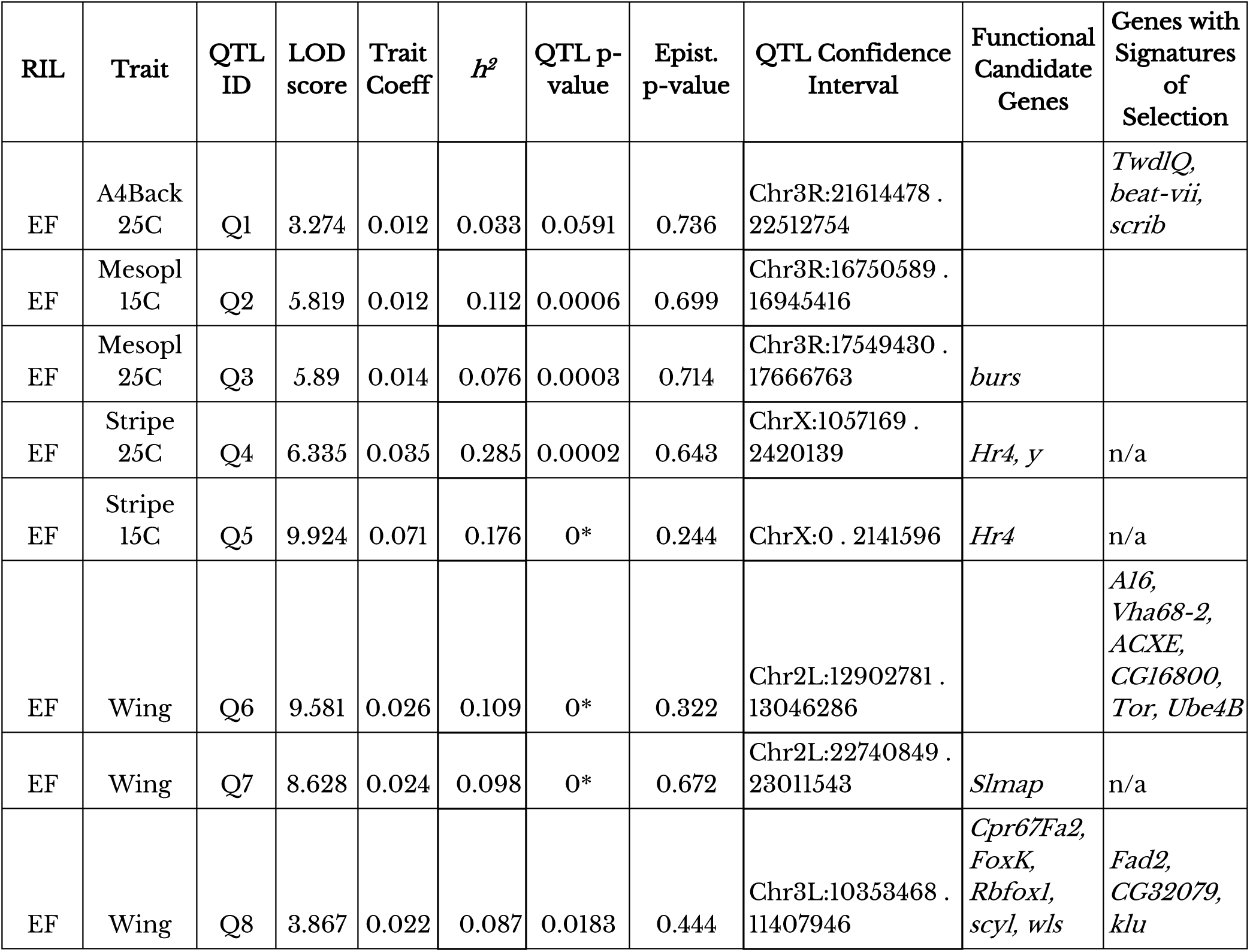
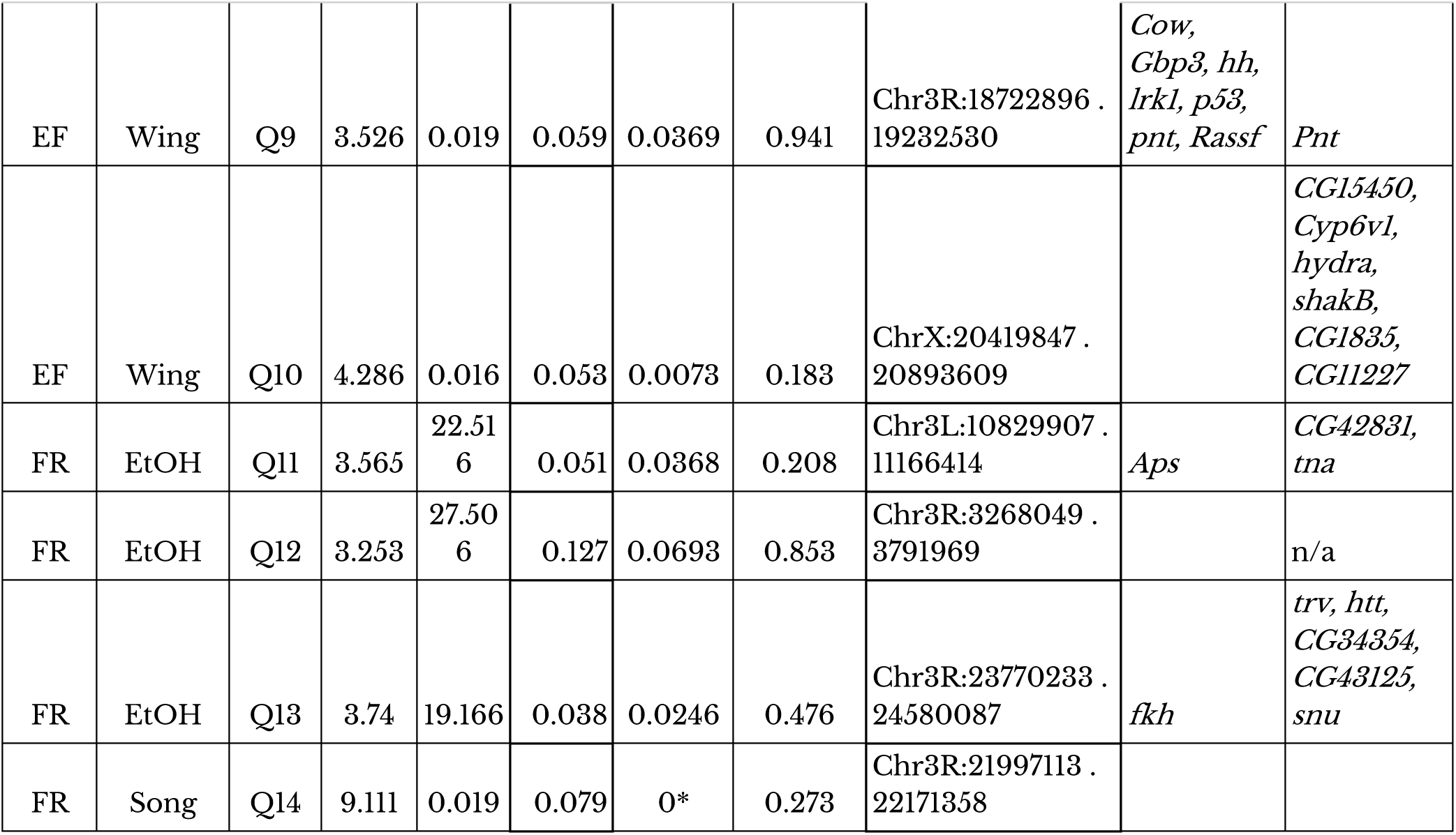
Characteristics of detected QTLs for all analyzed phenotypes, and possible underlying genes. The single QTL coefficient shows the magnitude of phenotypic change by substituting one allele of the derived population ancestry at that locus, which is shown on the scale of the measured phenotype. The heritability due to the QTL (*h^2^*) shows how much of the phenotypic variance is explained by the QTL (Broman *et al*. 2003). QTL p-value and epistasis p-value were obtained based on 10,000 permutations. * an estimated p-value of zero means the empirical value was more extreme than any permutation. Q4 was excluded from subsequent meta-analysis due to a stronger overlapping QTL for a correlated trait. Further details of each QTL are given in Table S8. Candidate genes are given in light of a priori functional knowledge based on FlyBase annotation or as population genetic outliers for local adaptation statistics (n/a indicates that QTL was in a region of low recombination and was not analyzed for signatures of selection). Genes that may underlie the song trait are instead described by Lollar *et al*. (2025).

Of the five pigmentation QTLs, the two identified for stripe ratio (Q4 and Q5, at 25 °C and at 15 °C respectively) overlapped. We mapped the residuals values of the correlation between these two traits to investigate whether there was a signal beyond their correlation and did not obtain any significant QTLs (Figure S7). Therefore, only the overlapping QTL with the higher LOD score (15 °C) was retained for meta-analysis (Table S8). Confidence intervals for the two QTLs detected for mesopleuron (one at 25 °C and one at 15 °C) were separated by just over 600 kb, so they were kept as two independent QTLs for the meta-analysis (Table 2).

The direction of all QTLs (Table 2) was in the expected direction: substituting alleles of the derived population resulted in phenotypic changes in the direction of the derived populations. This pattern is congruent with a scenario where these QTLs are contributing to locally adaptive trait changes (involving either the measured traits or else pleiotropically correlated traits under selection), given that if the phenotypic difference were neutral, we would expect some QTLs to show an effect in the opposite direction (Orr 1998).

### Outlier and Candidate Genes within QTLs

To generate hypotheses for possible causative genes underlying trait changes, we used population genetic summary statistics to identify genes within our QTLs that are potentially under local adaptation. Specifically, we flagged genes associated with windows that fell within the chromosome arm’s top 1% of windows for window *F_ST_*, maximum SNP *F_ST_*, or the haplotype statistic χ_MD_ (Table 2). These local adaptation scans were not conducted in low recombination QTL regions due to the expected lack of gene-scale resolution of selection signals in such genomic intervals. Since a striking signal of local adaptation is not guaranteed at loci underlying these strong population trait differences (*e.g.* if they involved soft partial sweeps), we also noted the presence of other functionally relevant candidate genes within each QTL.

We found a total of 26 candidate genes with functionally relevant annotations on FlyBase related to their specific measured phenotype. Q3 (mesopleuron 25 °C) included the gene *burs* (Dewey *et al*. 2004), implicated in cuticle tanning and hardening. Q4 and Q5 (the two stripe ratio QTLs)both included the candidate gene *Hr4*, a gene that when suppressed results in reduced pigmentation (Rogers *et al*. 2014). One also included the gene *yellow*, canonically implicated in pigmentation variation and evolution (e.g. Massey & Wittkopp 2015). This pair of QTLs overlapped others previously detected for the same trait in distinct Ethiopia-Zambia crosses (Bastide *et al*. 2016). The gene *ebony*, at which a soft sweep signal associated with a pigmentation QTL was previously characterized for Ethiopian *D. melanogaster* (Bastide *et al*. 2016), fell just between the closely-located Q2 and Q3 (mesopleuron 15 °C and 25 °C).

At least one gene related to wing development was found within each of the five wing length QTL regions. Within Q6 (on arm 2L), our population genetic outliers included the *mechanistic Target of rapamycin* gene (*mTor*), which plays a key role in insulin signaling and growth regulation, and has been tied to wing size and development specifically (Parker & Struhl 2015). Outliers inside this QTL also included the potential wing regulators *Vha68-2* and *Ube4B* (Krupp *et al*. 2005, Blanco *et al*. 2010, Okada *et al*. 2016, López-Varea *et al*. 2021). Within Q7 (low recombination regions of 2L), among the few genes present was *Slmap*, a hippo signaling gene shown to alter wing size (Zheng *et al*. 2017). Within Q8 (3L), population genetic outliers included *klu*, which regulates cell size and proliferation, and is associated with wing size defects (Schertel *et al*. 2015). Within Q9 (3R), outliers included *pnt*, related to wing morphogenesis (Dworkin & Gibson, 2006, Bejarano *et al*. 2008, Paul *et al*. 2013). The X-linked wing QTL (Q10) included an outlier gene possibly involved in wing development, *shakB* (Krishnan *et al*. 1993). Q7, Q8, and Q10 overlap with previously detected wing length QTLs, each from a different Ethiopia-Zambia cross out of four previously analyzed mapping crosses (Sprengelmeyer *et al*. 2022).

For the ethanol resistance QTLs, we identified two population genetic outlier genes (*htt* and *trv*) previously implicated in response to ethanol (Fochler *et al*. 2017) within Q13 (3R). Outlier genes detected within the other two QTLs had no annotated connection to ethanol. These QTLs did not overlap with those previously detected for the same trait from distinct France-Zambia crosses (Sprengelmeyer *et al*. 2021), in line with the genetically variable basis of this trait identified by that study.

Overall, we identified both novel candidate genes associated with the detected QTLs and genes that had already been implicated with their respective trait. The identification of genes already known to underlie these traits (although in many cases not their evolution) provides support for the approach we used here.

### Epistasis

To investigate whether there is evidence of epistasis among adaptive loci, we performed an epistasis scan for each single QTL and a meta-analysis combining the results across non-correlated traits. We did not identify any individually significant epistatic QTL pairs. The lowest genome-wide interaction LOD score p-value we obtained was 0.183, for a wing length QTL (Table 2).

Our meta-analysis of the combined non-overlapping single QTLs also supports the hypothesis that there is no strong epistasis involving adaptive loci. By combining the interaction p-values for all non-overlapping QTLs (where each p-value denotes the probability of that QTL having any interaction LOD effect that strong in the genome, based on 10,000 permutations), we obtained a p-value of 0.763 using Fisher’s combined probability test. This result indicated that the observed p-values do not deviate from the null expectation of uniformly distributed p-values expected without epistasis involving adaptive loci.

Because our sample sizes of RILs may not be suitable for detecting all magnitudes of epistasis, we conducted a simulation-driven analysis to indicate the strength of our negative results, and to indicate if there was a parameter space of epistasis that we could confidently rule out based on these results. Based on this analysis, our method had a high power (over 80%) to detect an epistatic QTL pair in scenarios of sign epistasis, in which the interaction effect changed the direction of the main effect, as well as high power to detect strong positive epistasis in which the interaction effected increased the main effect at least 2.5-fold (Figure 2, Table S9). We also tested the power of our meta-analysis by re-sampling 13 p-values from simulated data sets of different interaction effects (Figure 2). This analysis indicated that we should have had power to detect similar scenarios of strong epistasis as indicated for the lowest p-value analysis above (*i.e.* sign epistasis and positive epistasis at least doubling the main effect, and that we would likely have detected masking epistasis as well, in which the main effect was erased by the modifier). Therefore, although the power to detect moderate and weak epistasis is reduced rapidly, the lack of detected epistatic interactions from our empirical data supports the hypothesis that strong forms of epistasis did not play an important role in these instances of evolution.

**Figure 2.**
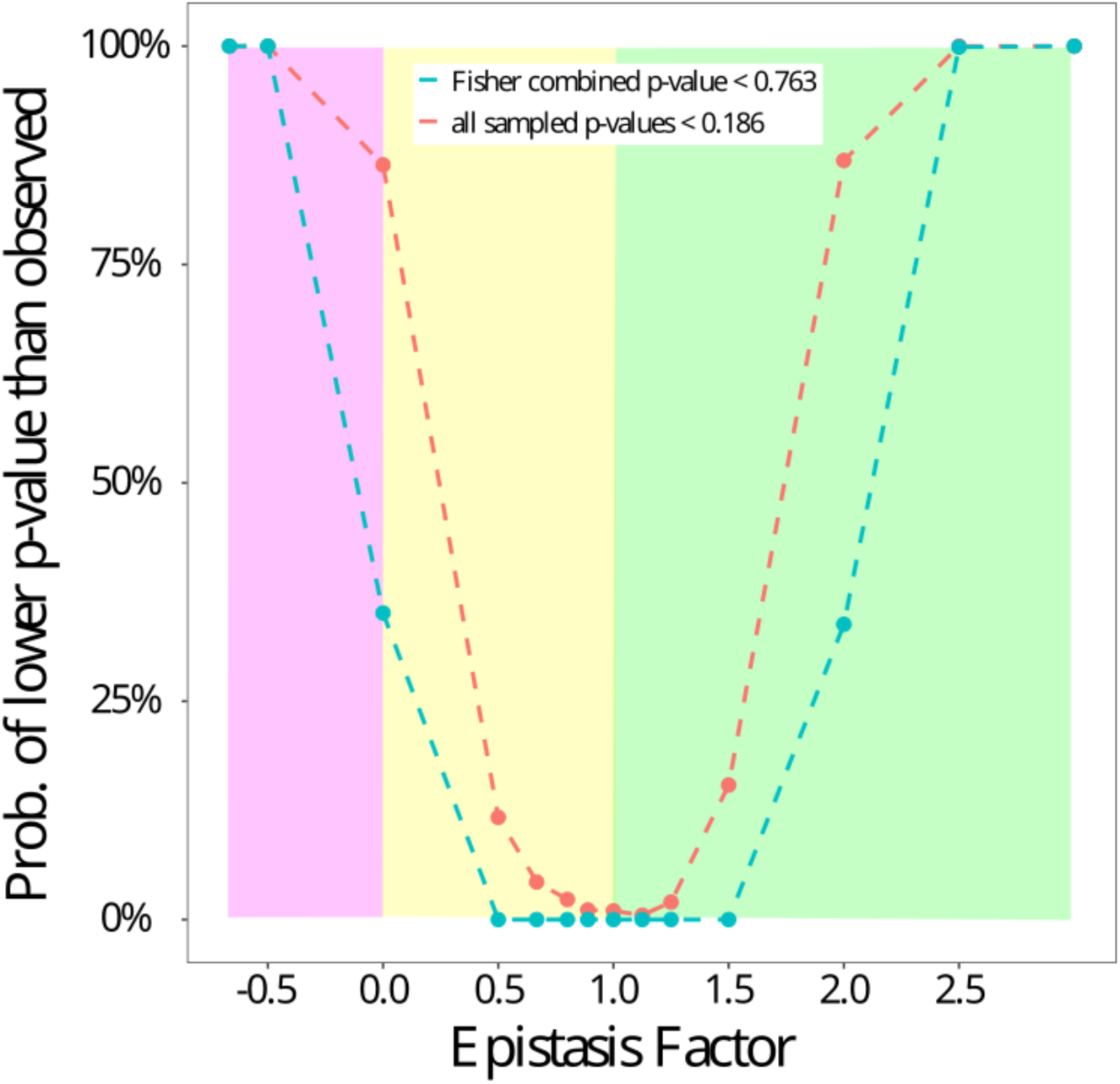
Simulation analysis indicating that strong positive epistasis or sign epistasis should have been detectable for the empirical data. Simulations based on randomly permuted individual genotypes and simulated epistatic effects (see Materials and Methods) were conducted to assess our ability to detect varying models of epistasis based on either the lowest empirical epistasis p-value (red series) or the Fisher-combined epistasis p-value across 13 analyzed QTLs (blue series). Here, the epistasis factor (x-axis) represents the multiplier that a modifier locus exerts on the primary QTL’s effect. Thus, negative values represent sign epistasis (pink shaded area), zero represents masking epistasis, values between 0 and 1 indicate negative epistasis (yellow shaded area), and values above 1 indicate positive epistasis (green shaded area). The y-axis shows the proportion of occurrences out of the 1,000 resampled instances in which a given model of epistasis yielded a lowest p-value lower than the observed 0.183 (red), or else how often the combined Fisher p-value was lower than the observed 0.763 (blue).

### Phenotype plasticity for pigmentation QTLs

Fly pigmentation is known for its temperature-based plasticity; flies reared at lower temperatures develop darker phenotypes than those raised at warm temperatures (David *et al*. 1990). Our results recapitulated this behavior, in that average pigmentation traits were higher (*i.e.* darker) at 15 °C (Figure 3). We identified one overlapping QTL for stripe ratio at both temperatures (Table 2). Based on the trait correlation and the shared QTL, at least part of the genetic architecture underlying adaptive melanism appears to be shared between temperatures.

**Figure 3.**
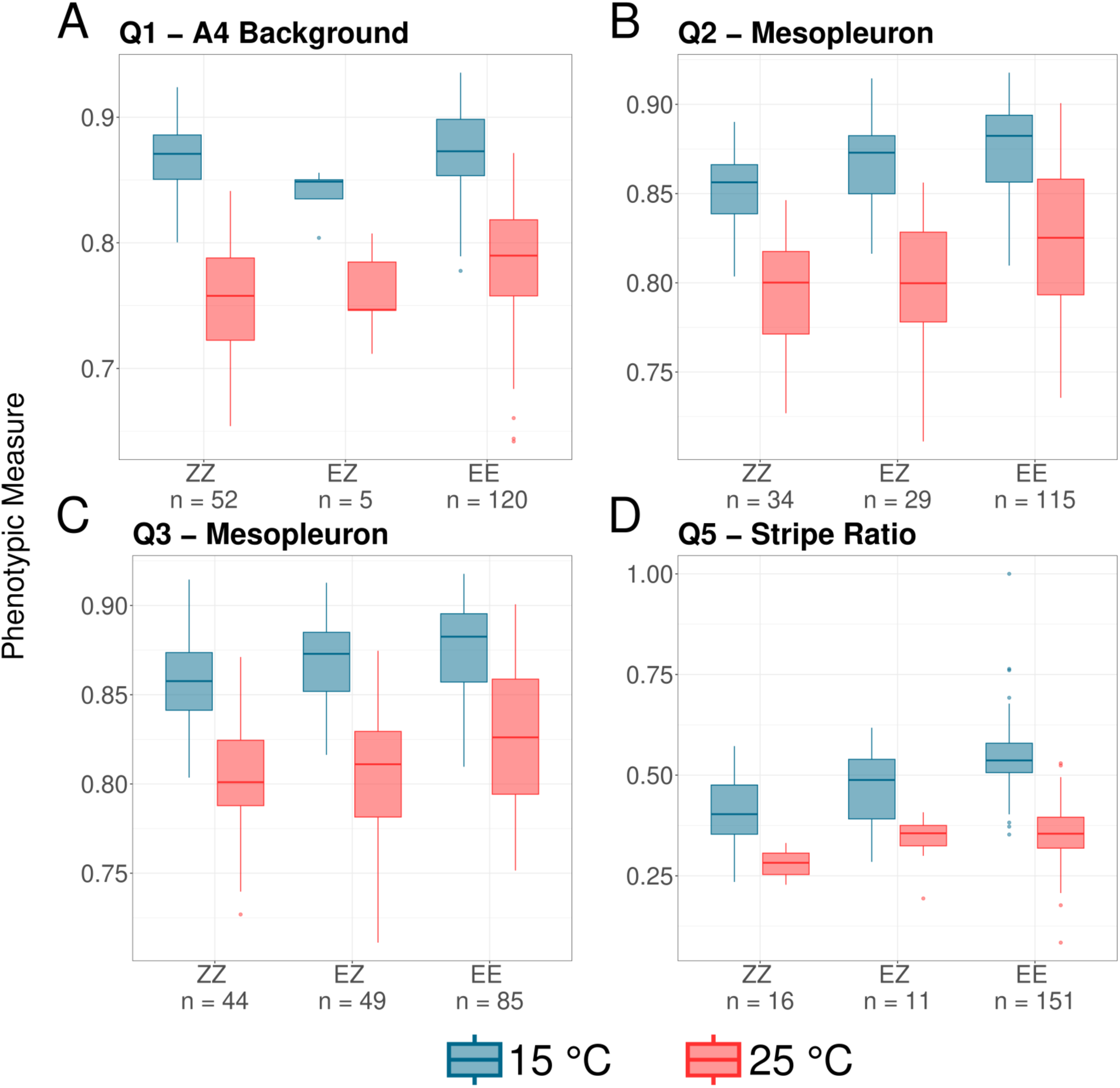
The reaction norms of the four non-overlapping pigmentation QTL show darker phenotypes for flies raised at 15 °C than at 25 °C and for flies with Ethiopia ancestry alleles, corroborating expectations. (A) Q1: A4 Background at 25 °C, with peak centered near 21.6 Mb on 3R. (B) Q2: Mesopleuron at 15 °C, near 16.8 Mb on 3R. (C) Q3: Mesopleuron at 25 °C, near 17.6 Mb on 3R. (D) Q5: Stripe ratio at 15 °C, near 1.3 Mb on the X. The y-axis shows the mean phenotype for each genotype and temperature treatment; a higher number is a darker phenotype. ZZ = Zambia homozygous, EZ = heterozygous, EE = Ethiopia homozygous.

The reaction norm of four pigmentation QTLs also showed that flies raised at 15 °C showed darker phenotypes for all traits and even darker phenotypes for the Ethiopia homozygotes, as expected (Figure 3). We also found significant genotype-by-environment interactions for two of these QTLs – A4 Background (p = 0.003) and stripe ratio (p = 0.007) (Figure 3A,D). The change in stripe ratio (Figure 3D) was greater for flies with the homozygous Ethiopia genotype. Here, the more derived-like environment appears to enhance the phenotypic consequences of the Ethiopian pigmentation variants. In contrast, for the A4 Background QTL, the phenotype means at 15 °C only show a small difference between homozygous genotypes, compared to a larger difference at 25 °C, which is congruent with this QTL being found only at 25 °C (whereas differences in the heterozygote means may reflect small sample size effects).

## Discussion

We report the creation of two new *Drosophila melanogaster* RIL panels established from crosses between single inbred lines from Ethiopia and Zambia, and from France and Zambia. We used these new data sets, together with an updated QTL mapping approach focusing on additive effects, to investigate the genetic architectures of multiple traits that have evolved to become differentiated between *D. melanogaster* populations. We did not find evidence for strong epistasis underlying their evolution, but genotype-by-environment interactions were supported for two of the three traits investigated.

### Genetic architectures underlying trait evolution

Our QTL mapping results indicate genetic architectures for most traits involving detectable loci with non-trivial effect sizes (5.3% to 21.4%; Table 2), consistent with past bulk mapping studies for some of these traits (Bastide *et al*. 2016, Sprengelmeyer *et al*. 2020, Sprengelmeyer *et al*. 2021). In combination with those studies, our results reinforce previous findings that evolved populations maintain persistent variability in the genetic basis of pigmentation traits, wing length, and ethanol resistance. We did not find significant QTLs for cold tolerance or thermal plasticity of pigmentation, which could reflect either a more complex genetic basis of those traits or else greater non-genetic variance in those assays.

We have also identified genes within the detected QTLs that might be underlying the relevant traits, including genes previously known to be related to the phenotype and, with the use of population genetic signatures of selection, novel candidate genes (Table 2). The identified genes represent viable hypotheses for contributors to the genetic architecture of traits that appear to have evolved due to local adaptation targeting either these or pleiotropically correlated traits but deserve further investigation. Additional studies using functional approaches such as genome editing will be needed to confirm the roles of these genes in the evolution of the respective traits.

### Lack of evidence for epistasis

Despite the pervasiveness of epistasis underlying the genetic basis of complex traits (Huang *et al*. 2012, Mackay 2014), we did not find evidence of epistasis involving any of the detected adaptive loci (Table 2). In light of our power analysis (Figure 2), our empirical results primarily argue against the presence of modifier variants within our RIL panels that trigger sign epistasis, complete masking epistasis, or strong positive epistasis with regard to the main effect loci. In contrast, our study does not speak to the presence of quantitatively negative epistasis (*e.g.* diminishing returns epistasis) or more moderate positive synergistic epistasis.

Previous studies have also found mixed results when investigating epistatic interaction among adaptive loci (Malmberg & Mauricio 2005). In some cases, the varied outcome for adaptive traits might reflect the transient nature of epistatic interactions in a population, given that the epistatic effects of two loci on a trait depend on the allele frequency of the interacting alleles. Interestingly, studies mapping QTLs for fitness-related traits (more directly related to reproduction and survival) have a more consistent result, often showing more epistatic QTLs than additive QTLs (Malmberg & Mauricio 2005).

One of the difficulties in extrapolating the results of experimental crosses to natural populations is that experimental crosses start with a limited amount of genetic variation (Mackay 2014). In our case, the crosses were made between one inbred line from each parental population, which will exclude some variants present in the source populations while elevating some naturally rare variants to moderate frequency. Previous genetic mapping studies of traits such as pigmentation (Bastide *et al*. 2016), ethanol resistance (Sprengelmeyer *et al*. 2021), and wing size (Sprengelmeyer *et al*. 2022) have shown that single crosses made with different inbred lines from the same population do not contain most of the genetic variation underlying population differentiation in each trait. However, our crosses have the advantage of focusing on a simplified genetic architecture, which may improve power to detect interactions among the variants present, in addition to clear inference of the parental origin of alleles genomewide.

We highlight that our study likely underestimates the epistatic interactions affecting *D. melanogaster* adaptive traits, since we only had power to detect relative strong interactions (Figure 2, Table S9). Based on the limited numbers of RILs available, we could only detect strong additive QTLs, and we needed to focus our epistasis scan on interactions with those few strong QTLs. Future research with a larger sample size (e.g. Torgeman & Zamir 2023) would be helpful to achieve more extensive insights into the scope of epistatic interactions affecting traits such as these. Studies using a more diverse set of starting lineages could mitigate some of the issues related to the limited genetic variation in our study. Additionally, drawing information from diverse sources, such as genome-wide association studies, bulk segregating approaches, and population genetics signatures of selection could be useful in designing mapping studies capable of not only identifying QTLs but candidate loci within QTLs. Multi-trait QTL mapping could also be a fruitful future direction to investigate the role of epistasis on adaptations as it can increase mapping power (Pitchers *et al*. 2019). Multi-trait approaches can also infer pleiotropy and epistasis (Zhang *et al*. 2016), adding a layer of complexity to the interpretation of results that could be especially challenging when some of the measured traits are not expected to have similar developmental and genetic bases.

### Genetic Architectures at Two Temperatures

Although the three body pigmentation traits we studied were somewhat correlated (Table S5), no shared QTL was noted between them (Table 2). Instead, we found two instances of overlapping or neighboring QTLs for the same trait measured at different temperatures, with 25 °C and 15 °C being more similar to the mean temperatures in the Zambian and Ethiopian populations, respectively. These results are compatible with the same or nearby genes being responsible for pigmentation differences at both temperatures.

*Drosophila* pigmentation is known to show thermal plasticity, and Ethiopian flies would have been expected to become somewhat darken on this basis alone, and yet they have also adapted genetically to become darker (Bastide *et al*. 2014). It is unclear whether this pigmentation plasticity is adaptive, which in itself can facilitate adaptive evolution (Ghalambor *et al*. 2007). Unlike larger ectotherms, it has been estimated that the solar gain in temperature from having dark pigmentation is at most a fraction of a degree for an organism with the small size and high surface to volume ratio of *Drosophila* (Willmer & Unwin 1981, Hirai & Kimura 1997). Across African populations of *D. melanogaster*, levels of ultraviolet radiation have been found to be stronger predictors of pigmentation than temperature (Bastide *et al*. 2014).

Gene-by-environment loci have recently been shown to be enriched for signatures of positive selection (Lea *et al*. 2022), supporting the hypothesis that environment-specific loci play an important role in local adaptation. Here, we did find that two QTLs underlying pigmentation evolution showed significant genotype-by-environment interactions (Figure 3), suggesting that the rate of their response to selective events might have changed upon colonization of colder environments. However, we did not find consistent patterns of either gene-by-gene or gene-by-environment interactions that met the predictions of cryptic variation. For example, the absence of epistatic interactions means that the Ethiopian alleles did not have their effect completely masked or reversed by another gene on the ancestral background. And in the case of gene-environment interactions, there was an enhanced QTL effect in the derived Ethiopia-like cool environment in just one of two significant cases (*i.e.* the stripe QTL). While cryptic variation may have played a role in the evolution of some or all of these traits, our results suggest that it may not have been pervasive with respect to the strongest effect changes.

## Conclusion

While the genetic architecture underlying adaptive evolution may vary somewhat depending upon the trait, there remains a key interest in assessing potentially general patterns. In combination with past studies, our results suggest that most adaptive trait changes involve variants of non-trivial effect size, and that these variants often do not initially reach fixation. Epistasis is an important component underlying complex traits, including in flies (Huang *et al*. 2012), but we did not find evidence that strong epistasis was relevant for the adaptive evolution of the studied loci. In light of the existence of other examples in which epistasis among adaptive loci have been shown (Malmberg & Mauricio 2005), our study highlights that the answer to the classic debate on whether epistasis is important to natural selection might vary case by case. Lastly, we found evidence of gene-by-environment interactions underlying pigmentation, stressing that the variability of genetic architectures can also be environment-dependent, which is particularly relevant in light of rapidly changing global environments.

## Data availability

Source code used in the analyses is available on GitHub (https://github.com/ribeirots/RILepi). Raw sequence read data for the Recombinant Inbred Lines are available on the Sequence Read Archive (BioProject: PRJNA861300).

## Acknowledgments

We thank members of the Pool lab for helpful comments on this manuscript.

## Conflict of interest

The authors declare no conflict of interest.

## Funding

This research was funded by NIH grants R01 GM127480 and R35 GM136306.

## Notes

### Competing Interest Statement

The authors have declared no competing interest.

### Summary of Updates

Various revisions after review, upon resubmission to same journal.

