## Supplementary material for "Recombinant inbred line panels inform the genetic architecture and interactions of adaptive traits in *Drosophila melanogaster*": Figure S

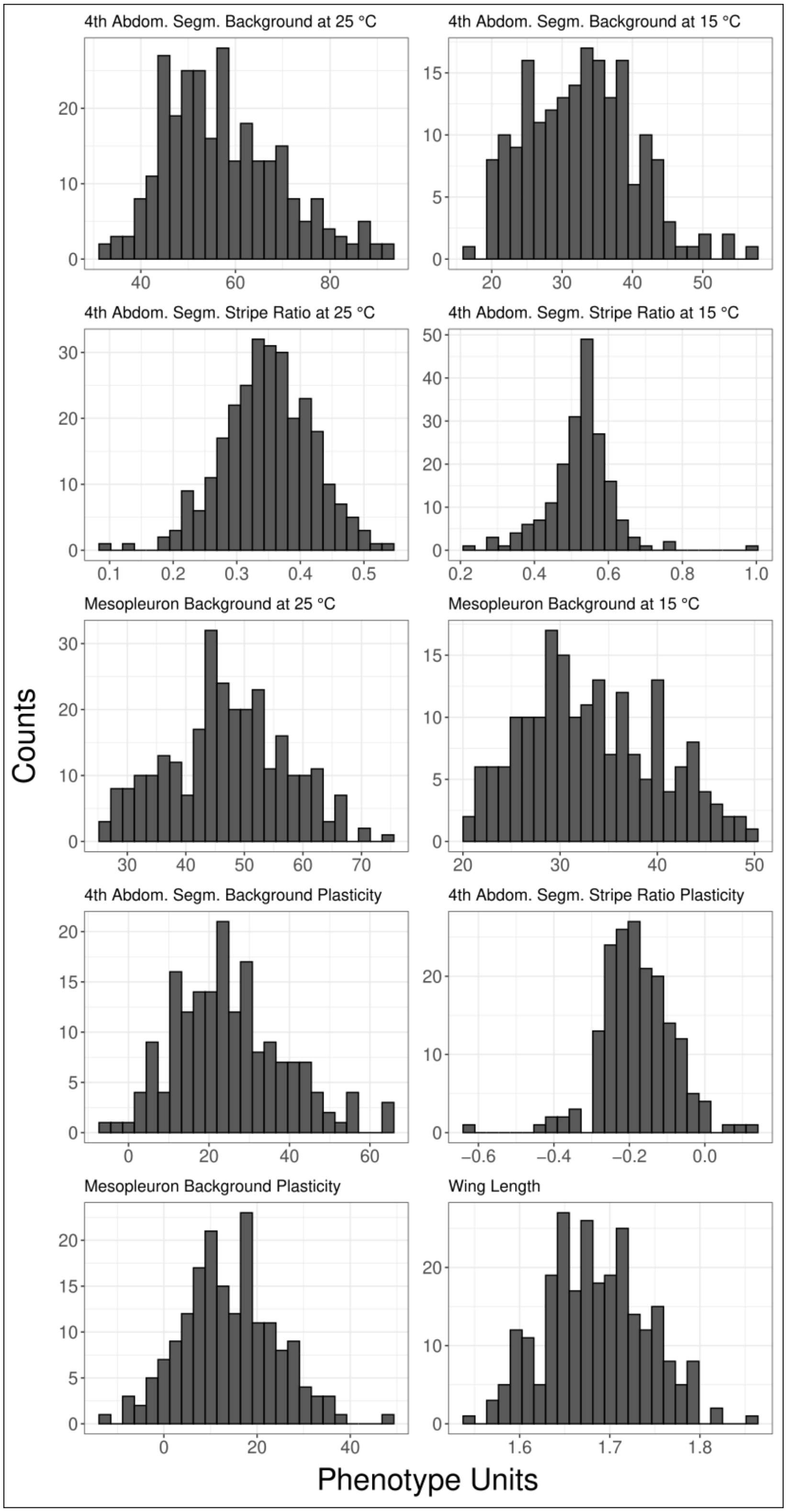


**Figure S1.** Phenotype distributions for each measured trait from the Ethiopian x Zambian RIL panel.

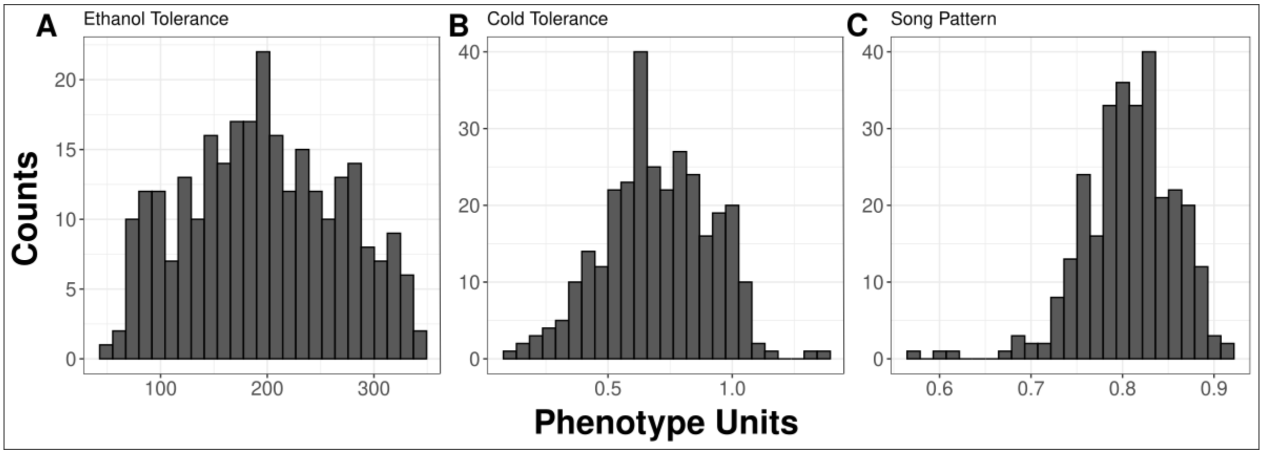


**Figure S2.** Phenotype distributions for each measured trait from the French x Zambian RIL panel.

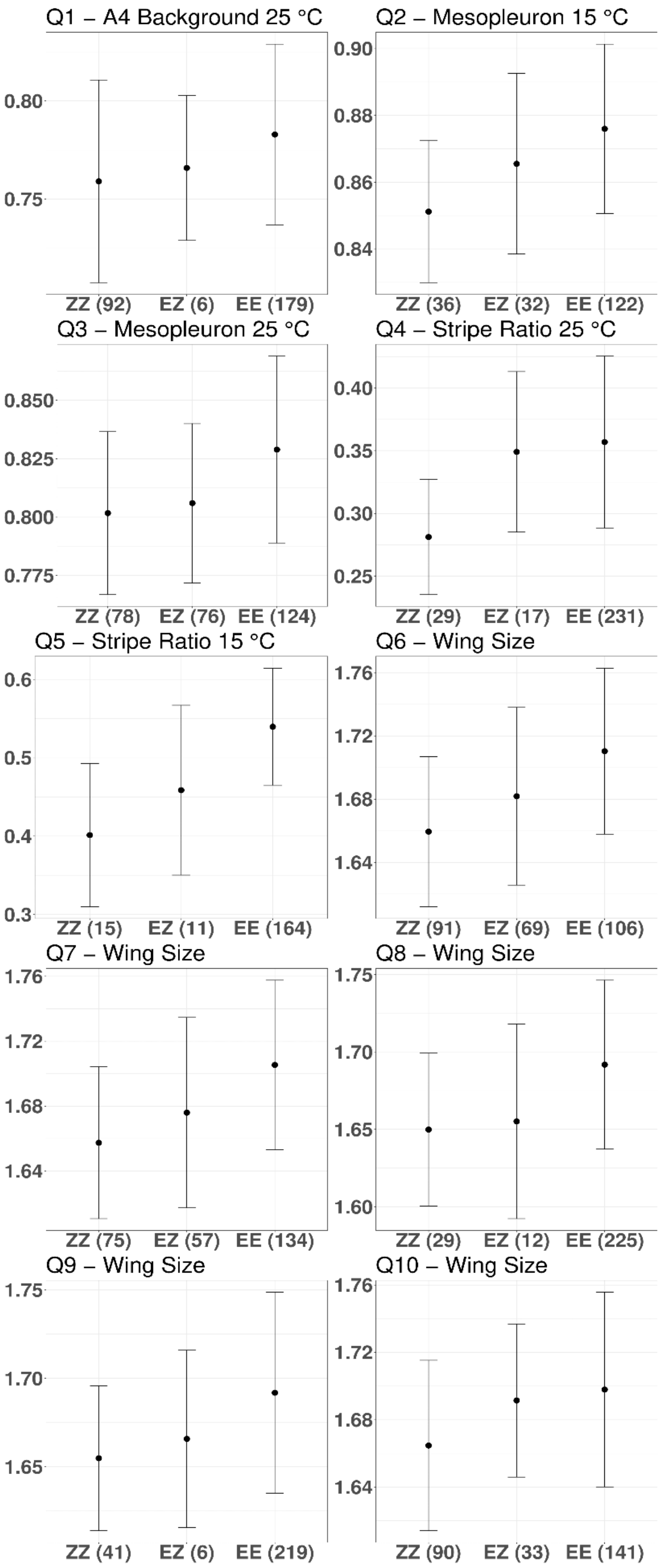


**Figure S3.** Phenotype by genotype distribution for detected QTLs from the Ethiopia x Zambia RIL panel.

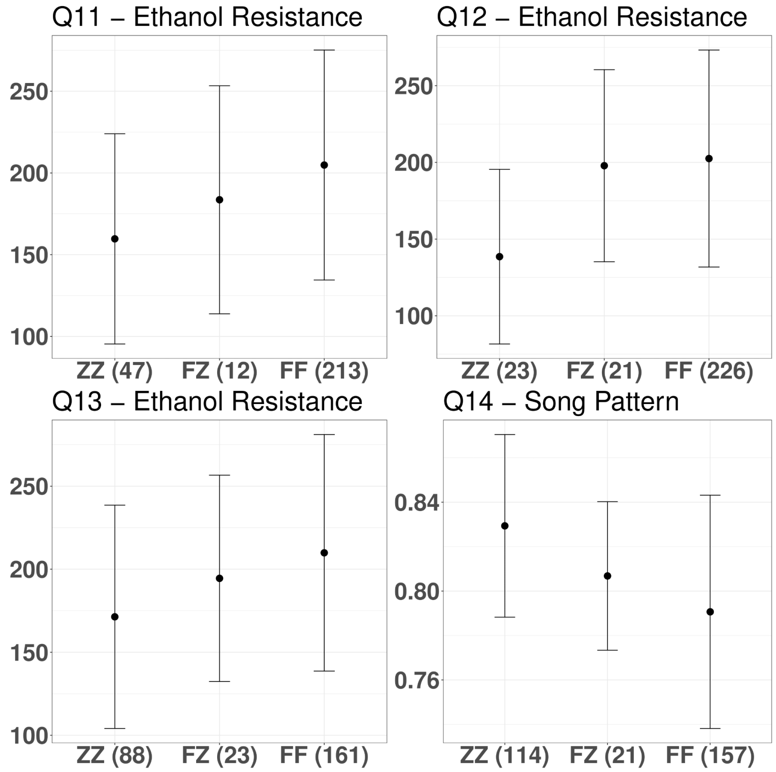


**Figure S4.** Phenotype by genotype distribution for detected QTLs from the France x Zambia RIL panel.

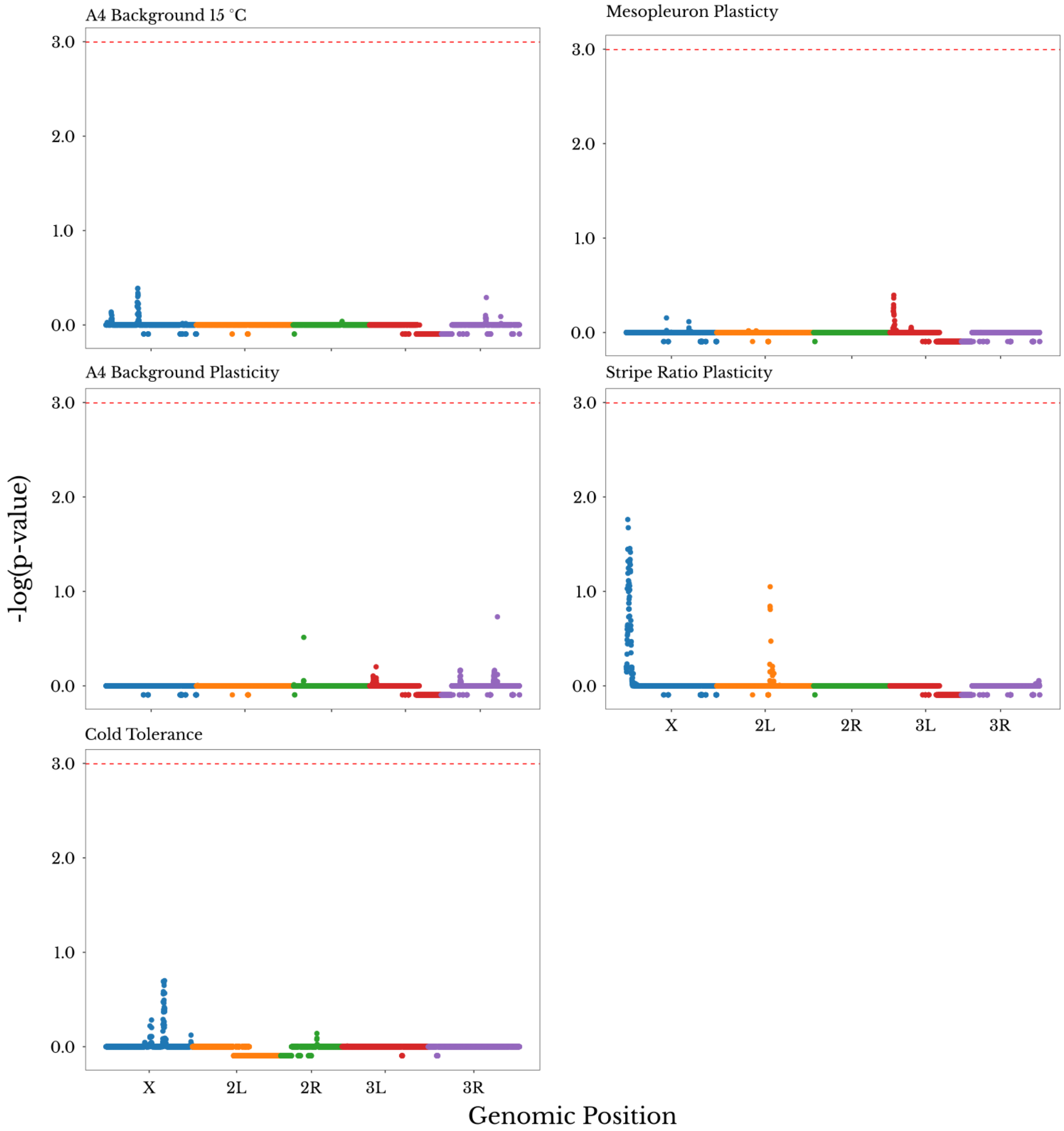


**Figure S5.** Genomewide LOD scores for traits without QTLs. Each panel shows the -log of the p-value for the LOD score of the genomic windows. Windows filtered out for ancestry skew are given a value of -0.25. The red dashed line represents the 0.05 p-value cutoff based on 10,000 permutations. The color of the dots represents the chromosome arm of each genomic window.

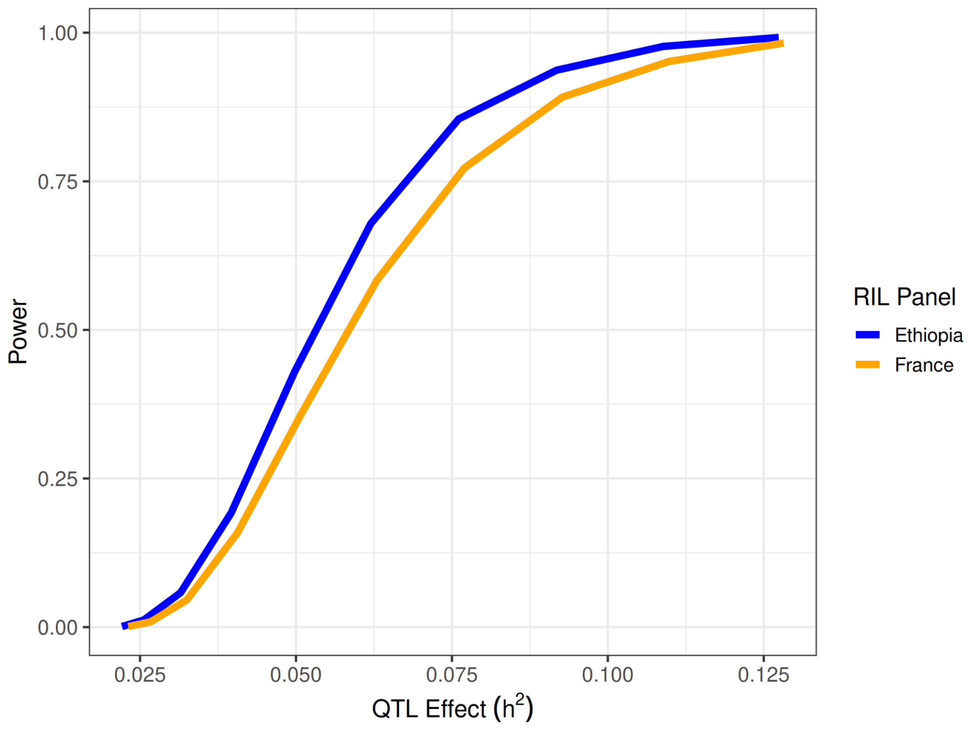


**Figure S6.** QTL mapping power for both RIL panels showed high power to detect QTLs of effect size 0.1 or higher.

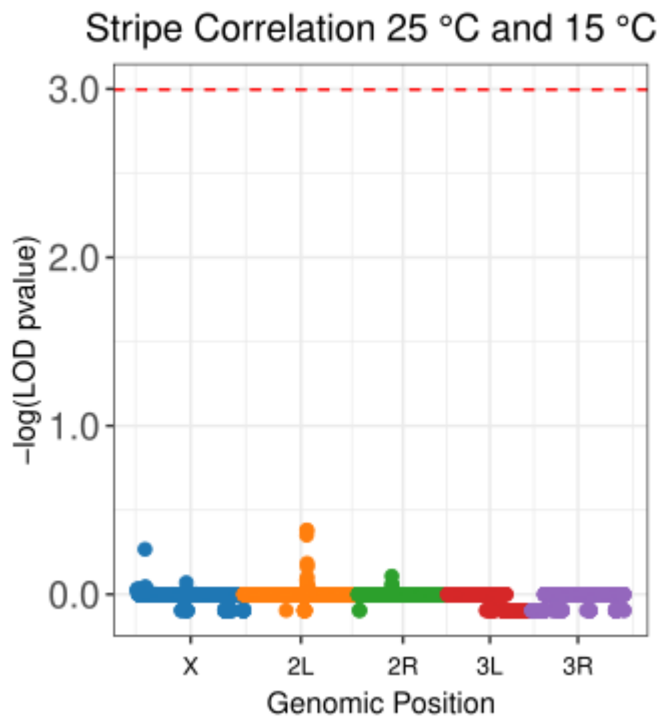


Figure S7. Genomewide LOD scores from the QTL mapping of the residuals of the two overlapping correlated traits (Q4 - stripe ratio at 25 °C, and Q5 - stripe ratio at 15 °C) showed no significant QTL peak – indicating a likely role of pleiotropy in the overlapping QTLs of this same trait at two different temperatures. Windows filtered out for ancestry skew are given a value of -0.25. The red dashed line represents the 0.05 p-value cutoff based on 10,000 permutations. The color of the dots represents the chromosome arm of each genomic window.
